# Single cell Edit Detection and Identification Tool (scEDIT): computational workflow for efficient and economical single cell analysis of CRISPR edited cells

**DOI:** 10.1101/2025.01.23.634562

**Authors:** Gajendra W Suryawanshi

## Abstract

Advances in CRISPR technology are revolutionizing gene therapy and drug discovery. The advent of single-cell DNA sequencing (scDNA-seq) is generating granular level insight into CRISPR induced genomic changes both intended and unintended. However, analysis of single cell data necessitates expensive high-performance computing (HPC) clusters or large data servers. Analysis of the single cell data is currently limited unaffordable computational cost and the lack of open-source software tools. To address this, we present scEDIT, a fast, lightweight, portable, and standalone software for pre- and post-processing CRISPR editing data from the Tapestri single-cell DNA-seq platform. scEDIT is memory-efficient, multithreaded, and compatible with most UNIX based systems. Tests using a low-cost desktop and public single cell CRISPR data demonstrate that the tool can efficiently process raw sequences, identify cell barcodes, count unedited and edited amplicons per cell, and outputs detailed filtered reads. Analysis of the single cell CRISPR data reveals indel patterns shared between in vitro experiments and unique indel profiles detected for in vivo study. Results further demonstrate the ability of single cell analysis in providing quantitative insights into the true zygosity of edited cell population. Although data shows a linear relation between indel frequencies by read count and cell count details of indel share between difference cells can only be truly explored with single cell data. Furthermore, scEDIT can analyze massively parallel base editing in human hematopoietic stem cells providing insights into the base editing patterns for hundreds of gRNAs targeting GATA1 and HbF gene. Importantly, scEDIT detected low frequency unintended and potentially harmful deletions introduced during the base editing. The reanalysis of two independent studies demonstrated the efficiency, stability, and portability of scEDIT making it an invaluable tool for uncovering new insights into the single cell data with limited and inexpensive computational resources.

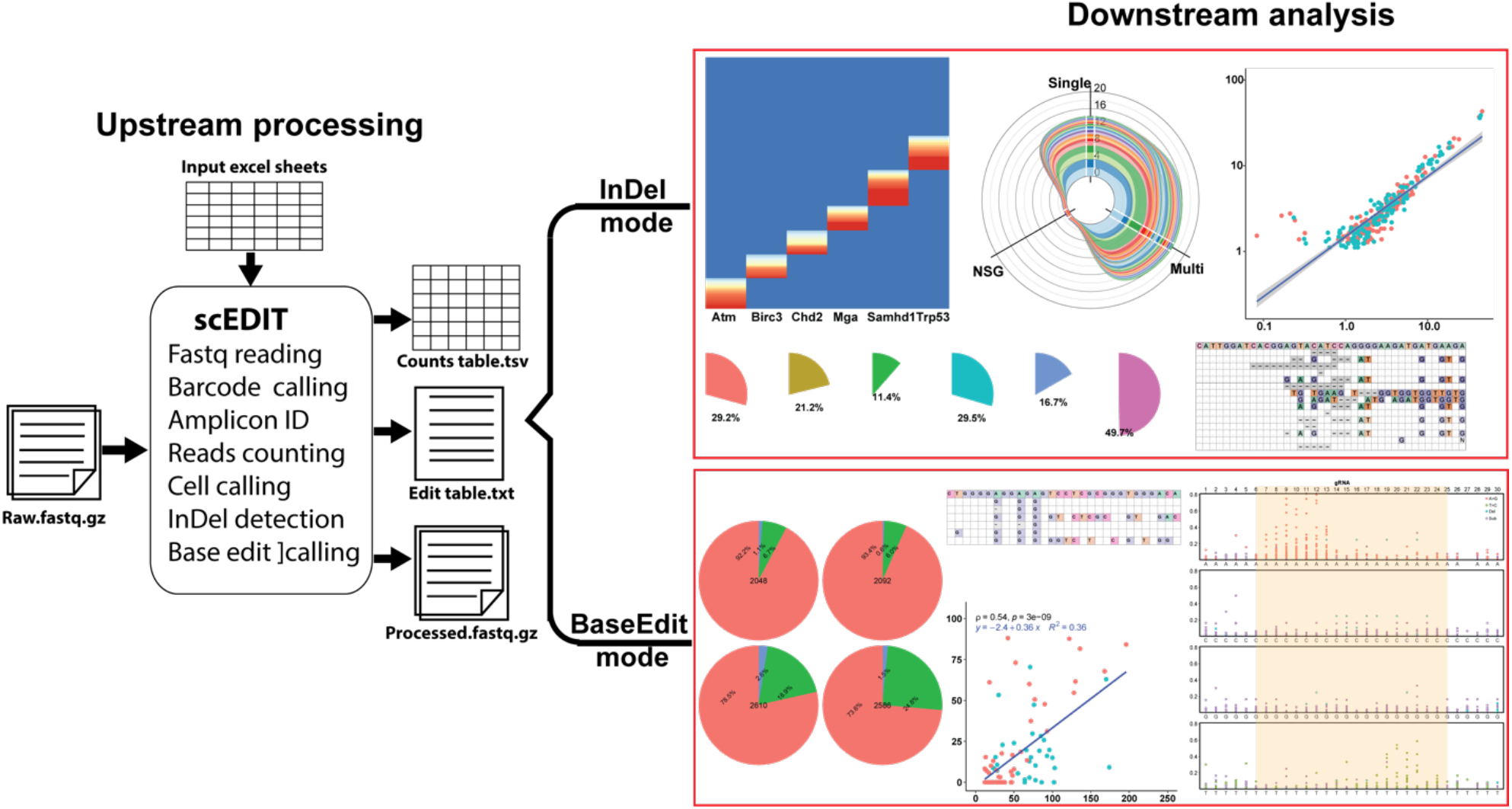

## Introduction

Advances in single cell sequencing are acerating biological research and revolutionizing the drug discovery space. Single cell multi-model analysis enables the definitive cell annotation and surface protein distribution across different cell subpopulations(*1*), multi-omics has provided single cell level resolution of epigenetic and transcriptomics landscape in different cell populations(*2*). These methods have provided valuable insight into cell type specific differences in gene profiles that are regulated by epigenetic modifications and transcriptome(*3*). Combination of single cell sequencing and CRISPR mediated precise gene silencing provides single cell resolution map of gene perturbation induced transcriptomics change and cellular response(*4, 5*). Analysis of DNA at single cell level has revealed the cell-to-cell heterogeneity of the mutation landscape in different cancers. Study used CRISPR-Cas9 to introduce mutations that disrupt gene function mimicking the loss of function (LOF) mutations observed in chronic lymphocytic leukemia (CLL)(*6*). The study used the single cell DNA-seq to track and quantify the CRISPR-Cas9 editing in the cells in vitro as well as in vivo and found bias in vivo growth in cells with indels in two specific genes. Advent of single cell analytics has enabled identification of link between rare yet detrimental genomic modifications and gene expression changes that are diluted in bulk cell experiments. The insights gained are transforming drug development and will be critical for the development of new cell and gene therapy strategies.

A recent study developed a massively parallel base-editing platform in clinically relevant primary human hematopoietic stem and progenitor cells (HSPCs) to systematically assess the functional effects of single-nucleotide variants during blood cell differentiation(*7*). Using combined pooled base editor libraries with single-cell RNA-seq and DNA-seq to map variant effects on cell fate and gene expression during hematopoiesis reveals the the causal roles of non-coding regulatory variants and missense changes in blood cell in health and disease. Edits were introduced by transducing the HSPCs with a pooled gRNA library, then the cells were electroporated with adenine base editors (ABE) to introduce A·T to G·C changes or cytosine base editors (CBE) for C·G to T·A changes. Cells were either maintained or pushed to differentiate into specific blood lineages (e.g. erythroid cells). Single-cell RNA-seq and separately pooled single-cell genotyping are performed to link genotype to cell fate and transcriptional profile. This study delivers a powerful, scalable approach to functionally profile genetic variants in primary human blood cells. Linking base edits to transcriptional and differentiation outcomes enables understanding of disease mechanisms, thereby improving the therapeutic interventions in blood disorders, and systematically characterizing clinically relevant genetic variants. While base editors are powerful and efficient, unintended small deletions are a real and measurable risk for therapeutic applications of BaseEditing. Thus, comprehensive profiling editing outcomes is critical and single-cell targeted genotyping can provide the granular view of the editing with zygosity and deletions at single cell level.

With FDA approvals of various viral vector-based gene therapies, the cell and gene therapies are becoming mainstream treatment strategies for cancer and rare genetic disorders. Despite success there are safety concerns arising for insertional mutagenesis caused by integration of vector in proximity of oncogenes. Recent clinical results of CRISPR based gene editing therapies are demonstrating the potential of gene editing and base editing approaches to cure diseases. However, the efficacy and off-target gene-editing remain a cause for concerns. Therefore, to critically assess the safety and efficacy there is need to develop single cell bioanalytical methods. Bulk cell analysis methods provide quantitative estimates for vector integration site analysis and for quantification potency and efficiency of CRISPR. However, the single cell resolution of single cell DNA sequencing can provide granular data for higher accuracy in quantifying true editing efficiency and zygosity resulting from partial editing.

Tapestri platform (MissionBio) provides single cell DNA-seq with targeted amplicon sequencing assay. The platform uses microfluidics system to partition the single cell in individual droplet and generate an emulsion of these droplets. Within the droplet cells are then lysed leaving behind genomic DNA and PCR reagents and barcoded primers. The PCR reactions take place within each droplet with genomic DNA of unique cells and generating amplicons with unique barcode for each single cell. The amplicons are then ligated with Illumina sequencing primers and libraries are sequenced paired end at appropriate depth. Raw data is then processed using proprietary Tapestri bioinformatics pipeline that performs cell barcode identification and generates a table of sequence counts for all the amplicons for each single cell assign with barcode. This matrix is then used for downstream analysis to determine valid cell based on the total sequence count as well as number of amplicons that were associated with the cell barcode. The pipeline provides automated cell calling, however depending on the study design manual thresholding approach can be more beneficial to improve the cell count. Currently, Tapestri bioinformatics pipeline is the only software tool available for pre and post processing of the Tapestri generated data. Access to the Tapestri bioinformatics pipeline is limited only to Tapestri equipment owners. Restricted access and expensive computational infrastructure requirements thus have prohibited reuse and reanalysis of one of its kind single cell DNA-seq data generated by Tapestri platform.

This study provides a novel software tool for analysis of Tapestri generated single cell DNA-seq data from CRISPR-Cas9 edited cells. Extending a previous computational workflow for detection of amplicons with LTR indexing in vector integration site sequences(*8–10*), single cell edit dictation and identification tool (scEDIT) provides a fast, lightweight, portable, and standalone software for pre- and post-processing single cell DNA-seq data of CRISPR-Cas9 edited cells. scEDIT is implemented in C++, Python and shell scripts, to make it fast and memory efficient and run on low-cost desktop thereby eliminating the requirement of high-performance computing (HPC) clusters. To demonstrate capabilities scEDIT single cell sequencing data from CRISPR-Cas9 edited cells that were expanded in vitro and in vivo was reanalyzed. Akin to Tapestri bioinformatics pipeline, scEDIT provides a matrix of cell barcodes and amplicon sequence count. Importantly, scEDIT is able detect short indels as well as large deletions in the amplicons introduced by CRISPR-Cas9 editing. Reanalysis of the data using scEDIT detected CRISPR introduced large (21 and 28bp) deletions in the clones that showed outgrowth in vivo. Further analysis also revealed the indel profiles and allele zygosity from in vitro and in vivo experiments, at single cell resolution. Reanalysis of the data demonstrated that scEDIT is an efficient alternative to Tapestri bioinformatics pipeline providing a tool for analysis of single cell DNA-seq data.

## Results

CRISPR-Cas9 gene editing provides a tool to introduce mutations to disrupt expression genes to study their role and function. Hacken et al(*6*) used CRISPR-Cas9 gene editing to introduce mutations that recapitulate the LOF in the 6 genes (TP53, ATM, CHD2, SAMHD1, MGA, and BIRC3) linked to CLL. Mutations in the genes were introduced either individually or in multiplexed fashion in murine interleukin 3 (IL-3)-dependent pro-B cells line Ba/F3. Tapestri single cell DNA sequencing platform was used to analyze the mutation induced in individual cells by single and multiplex gRNA CRISPR-Cas9. As a control experiment single cell sequencing was performed on add mix of 6 single LOF cell lines each of the six cell lines was edited separately for individual gene (here all referred as single edited cells. In addition, Hacken et al(*6*) also performed simultaneous edited of all 6 genes in the same cells (here on experiment is referred as multi edited cells). To further study the implication of LOF activity of the target genes in vivo the multi edited cells were infused in the NSG mice (here on referred as NSG experiment). Cells from all these experiments were independently processed with Tapesri platform that uses droplet-based single cell partitioning, PCR amplification of targeted genomics regions and NGS sequencing of the amplicons. Total 40 PCR primer-pairs were designed with different objectives: 1) 6 pairs to capture gRNA target regions in the 6 genes, 2) 22 for in silico predicted off-target regions, and 12 for internal positive control to improve cell calling. Hacken et al prepared the amplicon libraries for Illumina 2×150bp paired end sequencing and raw data from the sequencing processed using the Tapestri bioinformatics pipeline and CRISPR-Cas9 editing was analyzed using CRISPResso2.

### scEDIT requires minimal user interventions and outputs data in various formats

For reanalysis of Hacken et al data using scEDIT, raw sequence data was downloaded from the GEO as SRA files and converted to fastq file format using SRATools. Using the Trimmomatic (0.39) reads were quality trimmed and Illumina adaptors were removed. scEDIT was provided an excel file that contains metadata of the experiment with details of location of quality trimmed fastq files, version and location of reference genome used for primer and gRNA design, location of excel file detailing genomic coordinates of amplicons, excel file with gRNA sequence and CRISPR-Cas9 edit window with gRNA sequence with 20bp up or 20bp downstream. scEDIT converts the information to be provided as input for file processing of the data. scEDIT performs the following task for each paired read isolation and identification of cell barcodes detected in forward reads, amplicon sequences detected in both reads, scan read for CRISPR-Cas9 edits in edit window provided as metadata. Reads shorter than 25bp, missing cell barcode, missing or mismatched amplicons are filtered. Names of the reads that passed the filtering steps were updated by adding the cell barcode sequence, amplicon name, and edits status with short CIGAR text. scEDIT simultaneously counts reads all the amplicons associated with each detected cell barcode and for amplicons designed to capture CRISPR-Cas9 on and off-target editing records number of edited and unedited reads. scEDIT output the results in multiple different formats. All valid reads are stored in fastq format with read name containing cell barcode, amplicon name, edit status of the read. In these fastq files, forward reads are trimmed off the Tapestri barcode and constant sequences. The fastq files can be directly used as input files for other alignment tools. Matrix file with cell barcodes as rows and amplicon counts in columns for all the cells with valid barcode. scEDIT also outputs text file with list of all cell barcode and number of edited reads linked to the barcode. Additional package can be used to generate text files with columns cell barcode, read name, amplicon name (only gRNA on-target amplicons), edit status, and sequence with deletions represented as ‘-’ for better visualization of the CRISPR-Cas9 induced deletion.

### scEDIT provides efficient sequence processing at low computational cost

scEDIT is developed on low-cost desktop computer with objective to make processing and analysis of Tapestri data cost and time efficient. The low-cost desktop configuration includes low-end AMD Ryzen 7 5700G (8 core/16 thread) with base clock speed of 3.8 GHz processor, 128 GB of memory (RAM), and sufficient data storage space for large Tapestri data sets. To increase the read processing throughput raw sequencing data processing applications are implemented in C++ with ability for parallel processing for optimal use of multi-threaded CPUs. To lower the memory requirement scEDIT splits the raw sequence files prior to processing and final output is merged results from all the split files. This approach reduces the bulk memory requirement to process large single cell sequencing data files. scEDIT utilizes highly efficient C/C++ implementations of Smith-Waterman and Wavefront algorithms for detection of cell barcodes, amplicon sequences, and CRISPR-Cas9 edits. scEDIT used Python script as wrapper to run the entire analysis workflow from raw sequence processing to cell calling. Use of low overhead C/C++ programming enables fast, parallel, and memory efficient data processing that also made scEDIT both lightweight and portable. To demonstrate efficiency and accuracy of scEDIT, we processed the raw single cell data from Cas9 editing study of Hacken et al(*6*) and base editing experiments of Martin-Rufino et al(*7*)

Hacken et al(*6*) provided data for three experiments each has pair of fastq files with forward and reverse reads with approximately 15 million reads in each file. The desktop computer scEDIT was able to process ∼1 million paired reads per min using 6 cores or 12 threads totaling ∼15 min per sample/experiment. This process includes reading input fastq files, processing reads, barcodes and amplicon counting, detection of possible edits, writing multiple output files, and performing cell calling analysis. The time required to complete the read processing from raw to counts matrix and edit status of the read is fast enough to complete processing of multiple samples on the desktop and we also observed significant improvement in processing speed with more processing cores. Since scEDIT splits large raw sequence files in smaller files with lesser reads increasing memory capacity is not critical to speed improvement. These results show that scEDIT is both cost and time efficient tool for analysis of Tapestri data.

### scEDIT accurately identifies cell barcodes and amplicon primer sequences

Accurate identification of barcodes, constant, and amplicon sequences is critical for determining the true cell and amplicon counts. Accuracy of scEDIT in filtering out ambiguous barcode combinations or amplicon sequences is assessed using in silico generated sequencing reads. scEDIT identified the reads having valid barcodes with ∼100% accuracy, barcodes combination with 1 mismatch with 87.7% accuracy, and barcodes combination with 2 mismatch with ∼77.9% accuracy. scEDIT successfully removed ∼100% of reads with incorrect amplicon pairs. For in silico data composed of reads containing no error, barcode with 1 or 2 bp errors, amplicon mismatches, and/or errors in the sequence, the barcode recovery is up to 95% (supplementary figure s1). Stringency of sequence matching can be lowered or increased by respectively decreasing or increasing the sequence identity parameter. Since 9bp barcode with more than >2 mismatch leads to barcode conflicts, lowering the identity value causes misleading read assignment. To minimize the errors, a default value of max 2 mismatches per barcode combination (1 per 9bp barcode) is used. However, in case of highly similar amplicon primer sequences increasing sequence identity value can reduce the errors in identification of amplicons. Overall, the analysis of in silico data shows that scEDIT can accurately identify barcodes and amplicon accurately and efficiently.

For processing raw sequence data from Hacken et al(*6*) study barcode and constant sequence identity values are set to 90% and identity of amplicon primer sequences is set to 90%. However, due to high similarity between Trp53 (TP53) on and off target amplicon primers, identity value was set at 100% to avoid amplicon conflict.

### scEDIT can detect CRISPR-Cas9 editing and heterozygosity at single cell level

Bulk sequencing of the three experiments showed the level of editing in each experiment providing an overall view of the editing efficiency. Tapestri platform provides single cell targeted DNA-sequencing data and aims here is to provide a software tool to process this data efficiently and at low computational cost. To demonstrate capabilities of scEDIT, the single cell DNA sequencing data from these three experiments was processed with scEDIT run on a low-cost multicore desktop computer. Cell barcodes with more than 750 total sequence counts were considered as true cells and used for further analysis. For single edit experiment, each cell line was edited impendent of for one gene thus each cell line should only be edited at one gRNA target locus or its associated off-target sites, therefore cells with edits in more than one gRNA target locus are considered as possible doublets during the Tapestri platform. Using total read count cutoff scEDIT identified 3,543 single cells for single edit experiment, 3,151 cells for multi edit experiment, and for sample from NSG mice 2,517 cells. After filtering out ambiguous editing (the possible doublets) the final single cell count for single edit experiment became 3,304 cells. The editing status of these individual cells was determined based on the percentage of reads with edited sequence. Cells with more than 20% edited reads were counted as edited cells (Figure 1a, Heatmaps). In three edit experiments, the efficiency of editing at each target location (Figure 1b, Pie charts) is markedly different. In multi edit experiments, the percentage of editing is different from that observed in single edit experiment. The editing percentages observed in the NSG experiment are uniquely different from both single edit and multi edit experiments, likely due to in vivo growth advantage to the cells with editing in specific combination of genes.

**Figure 1.**
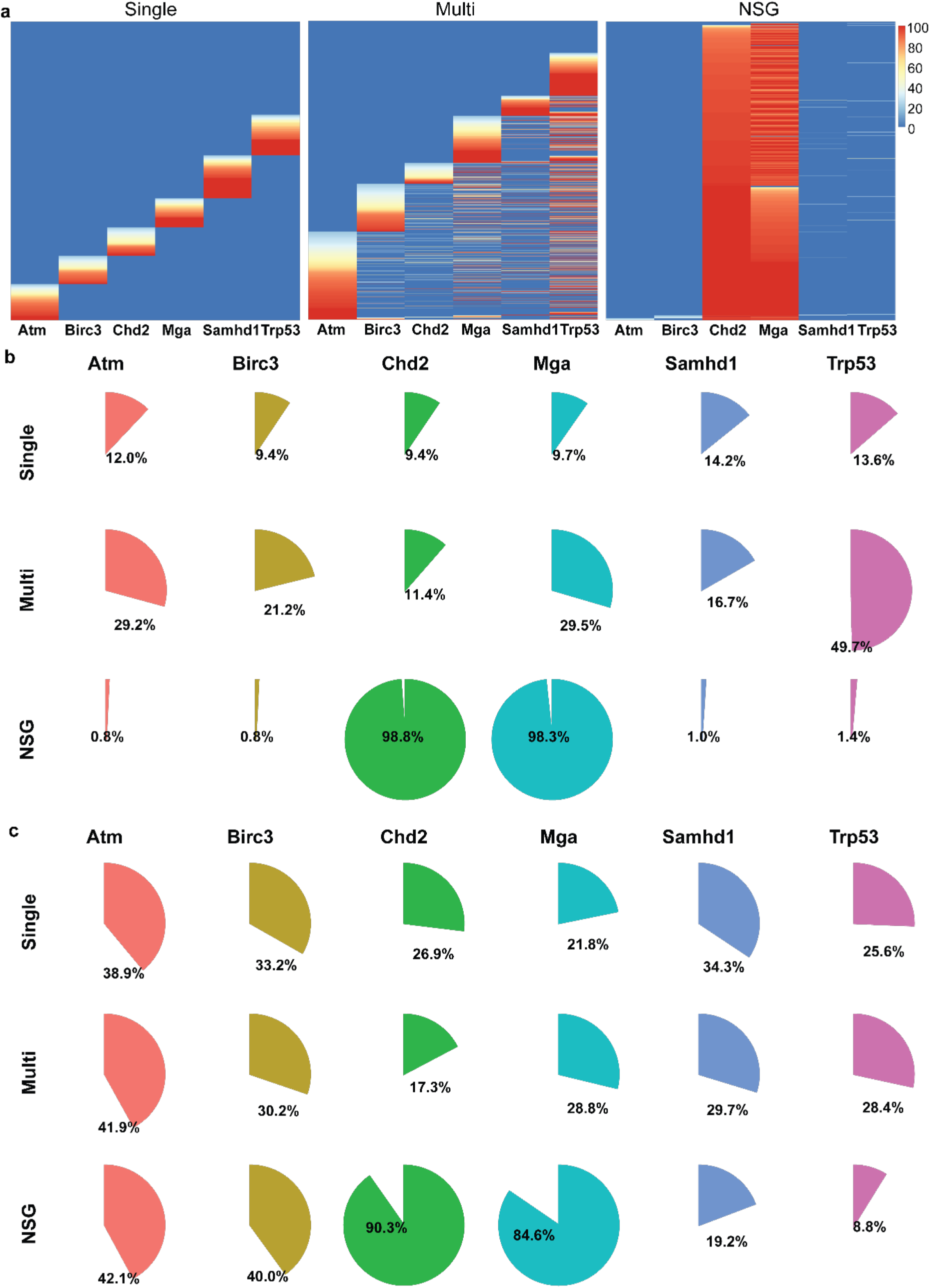
scEDIT identifies edited cell population in three different experiments. a) Heatmap showing edited cells in three different experiments each column in the heat map represents target loci and each row is one cell (cell barcode). Level of editing is on the scale of 100 to 0% based on percentage of edited reads in the cell (<20% edited reads is considered no editing i.e. set to 0). b) Pi charts showing percentage of edited cell for each target locus in three experiments. c) Pi charts showing percentage of cells (within the edited population) with heterozygous editing.

Taking advantage of the ability to analyze single cell indel profiles, scEDIT can distinguish between edits at different allele in the same cells. Cell having both unedited and edited reads or edited reads showing more than one indel pattern (both >20%) is indicative of different editing at two alleles or heterozygous editing. Cell with edited read percentage >80% for one edit is considered as both alleles having same indel pattern or homozygous editing. This editing cut off enables the quantification of hetro and homozygously edited cell populations. Analysis of scEDIT processed data shows different levels of zygosity for each of the six target genes and is different in each of the three experiments (Figure 1c, Pie charts). These results demonstrate the functionality of scEDIT in detecting and identifying CRISPR induced edits at single cell level and the ability to determine the zygosity of the CRISPR mutation in individual cells. scEDIT can also be used for deeper analysis for the CRISPR induced indel profiles and their distribution across the edited cell population.

#### scEDIT identified common indel patterns across different cells

Analysis of bulk cell data can provide the overall frequency of indel profiles in a population of cells. Single cell analysis of editing patterns using scEDIT can reveal specific indel profiles in each individual cell for each of the six target loci. Allele level detection of the edit patterns in individual cells enables quantification of the most recurrent homozygous and heterozygous indel pattern in edited cell population. scEDIT generated data thus allows identifications of common indel patterns and allele combination for each target loci across all the edited cells.

For single edit experiment, scEDIT detected indel patterns in each edited cell (Supplementary figures s2). The highest frequency indel sequence of each target locus is present in majority of the cells (Figure 2a) and was also detected in heterozygous cells. Data showed the prevalence of common indel in multiple cells with some indel appearing in pair in heterozygous editing (Figure 2b). Since each target was edited independently the high frequence indel sequences did not show up in cell edited for other target loci. Although some cells did indicate the presence of more than one target edit, these cells were very rare and the presence of such multi target editing likely resulted from PCR or sequencing errors in cell barcodes. In contrast, data from multi target editing experiments showed simultaneous editing at multiple target loci in many cells. Data also revealed the presence of common indel sequences between different cells. Like single edit experiment, the most frequent indel sequences (Figure 3a) were detected in the highest number of cells. However, in contrast to single edit experiment, in multi edit experiment the frequent indel sequences for each target loci (Figure 3b) were also detected in cells with editing identified at more than one target loci (Figure 3b). In this analysis shows that scEDIT can find the common or highly frequent indel sequences and quantify its presence in the population of edited cells. This analysis reveals the allele level to cell population level indel profiles and identified common indels across the cell population. Furthermore, analysis indicates presence of preference for indels across different experiments.

**Figure 2.**
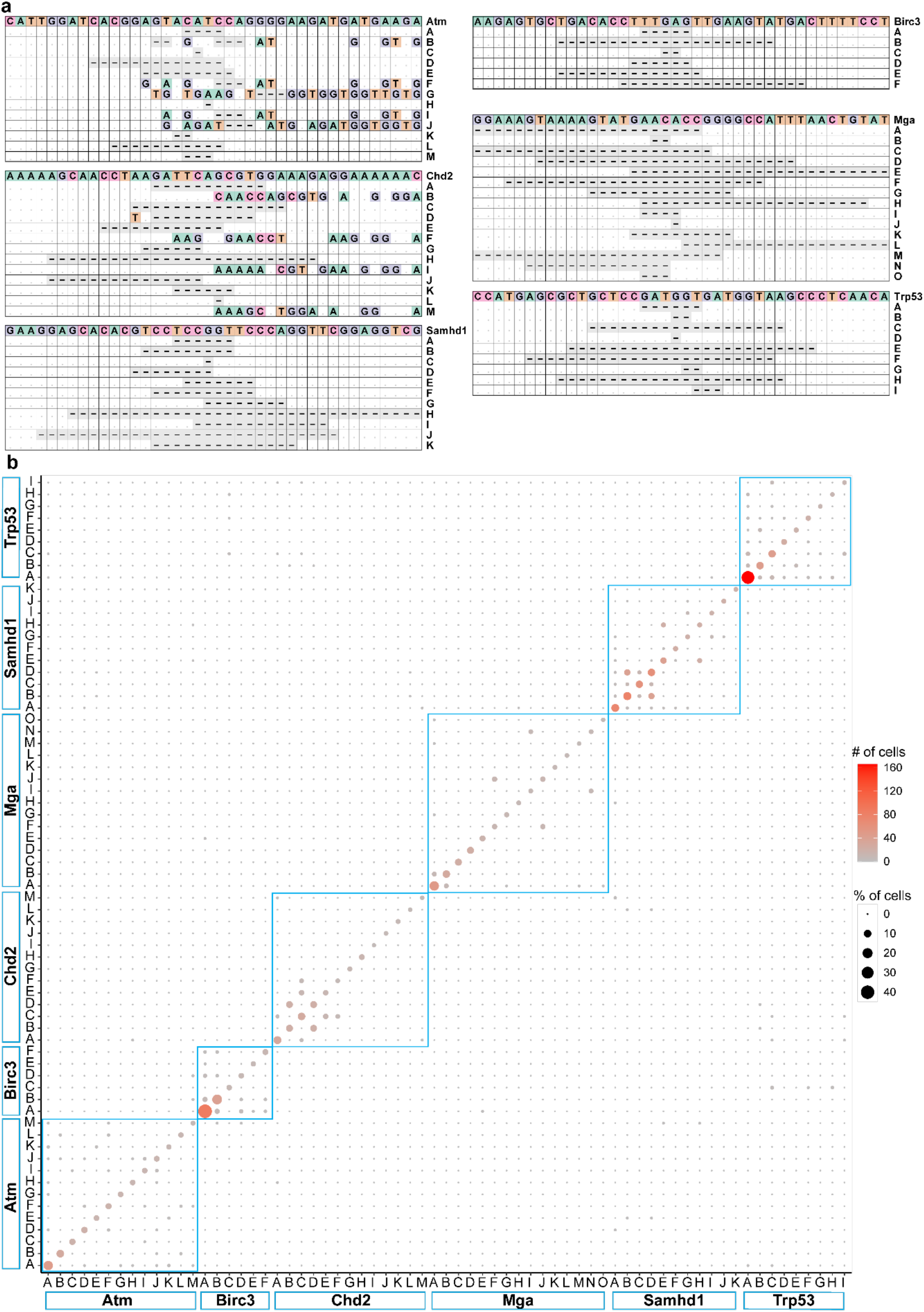
Indel profiles and their prevalence across multiple cells in single edit experiment. a) Indel profiles of 6 target loci and for each loci sequences ordered (top to bottom) by abundance (frequency) of the indel sequence in the total reads of the target loci. In indel profile plots the topmost sequence is reference sequence, mismatched bases are shown with respective nucleotides and deletion, and match are shown using ‘-’ and ‘.’, respectively. b) Bubble plot hosing the prevalence of indel high frequency indel in different cells and their pairing with other indel sequences. Size of the bubble corresponds to % of cells that carry the indel and darker color corresponds to higher absolute number of cells.

**Figure 3.**
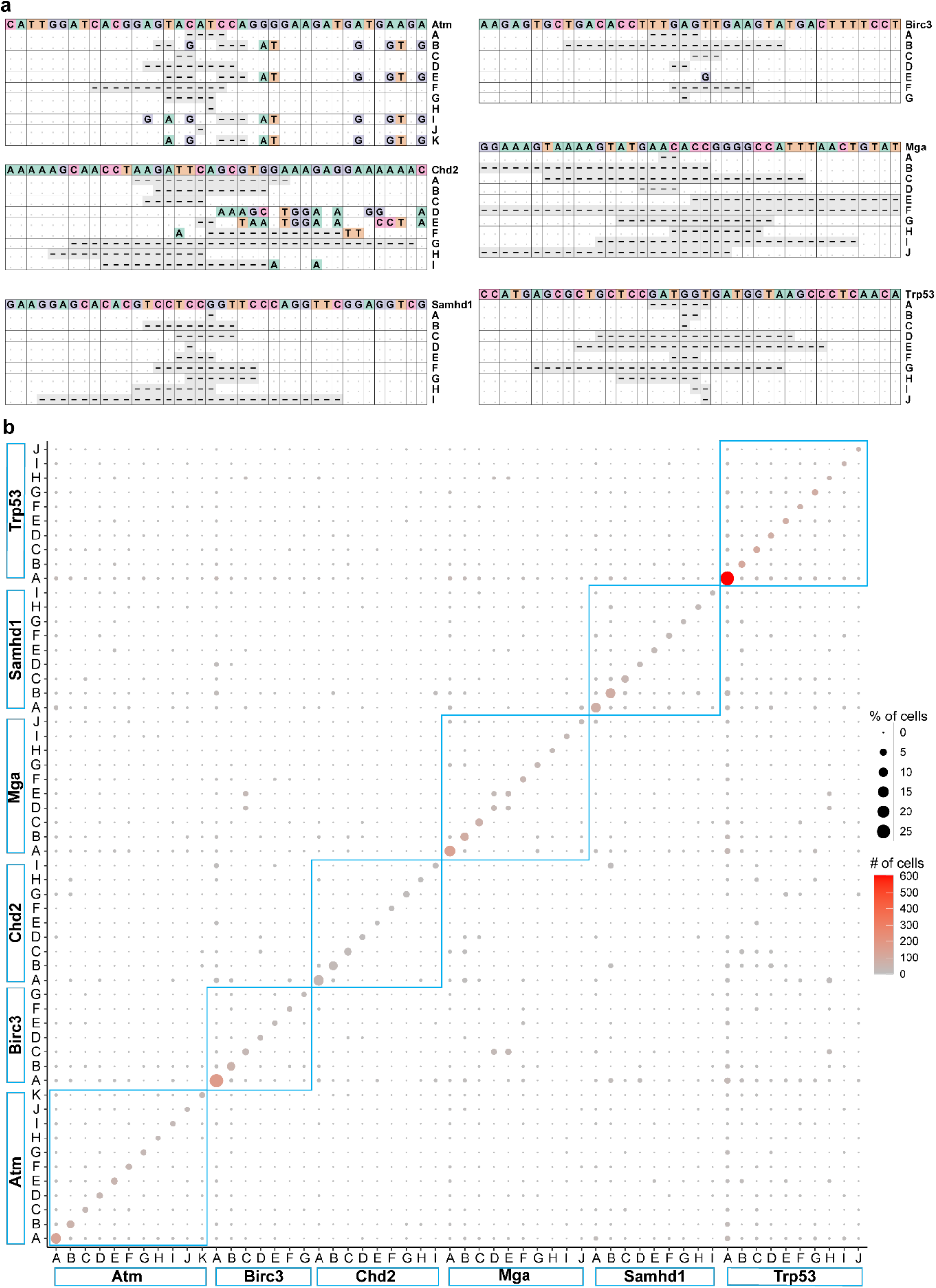
Prevalence of indel profiles across multiple cells and multiple target loci after simultaneous editing in multi edit experiment. a) Indel profiles of 6 target loci and for each loci sequences ordered (top to bottom) by abundance (frequency) of the indel sequence in the total reads of the target loci. In indel profile plots the topmost sequence is reference sequence, mismatched bases are shown with respective nucleotides and deletion, and match are shown using ‘-’ and ‘.’, respectively. b) Bubble plot hosing the prevalence of indel high frequency indel in different cells and their pairing with other indel sequences. Size of the bubble corresponds to % of cells that carry the indel and darker color corresponds to higher absolute number of cells.

### Quantification of indel overrepresentation in vitro and in vivo

The goal of introducing mutations in six target loci was to mimic the somatic mutations and explore the implication such mutations on cell growth(*6*). Analysis shows that for all the target loci, certain indel sequences were found in both single and multi-edit experiments at relatively similar abundance in the edited cell population (Figure 4). Similar indel profiles in both in vitro experiments, single and multi-edit experiments, indicate that certain indels are likely to have improved the growth potential of the edited cells. However, analysis of NSG experiment (in vivo experiment) revealed distinctively different indel profiles. Single cell analysis cells isolated from NSG mice showed that >97% of cells had edits in two loci, Chd2 and Mga, with very few cells having edits at any other 4 loci. scEDIT found two unique indel sequences for both Chd2 and Mga loci reflecting heterozygous editing of the cells (Figure 4, Chd2 and Mga petal plots). Despite the overrepresentation of Chd2-Mga edited cells in vivo, the indel sequences were not detected in vitro. Even for other 4 loci, indel profiles were specific to in vivo expansion and not found in in vitro (Figure 4). scEDIT has been able to both detect indels and quantify their abundance in cell populations. This enables a quantitative comparison of indel and its possible impact on cell growth. Although bulk cell analysis provides quantification of indel based of reads detected among sequenced reads information about the zygosity of the cell population remains unclear. Furthermore, the relation between abundance of indels by read count and by cell count need to be explored to validate the interpretation of read frequence based quantification of the bulk cell data.

**Figure 4.**
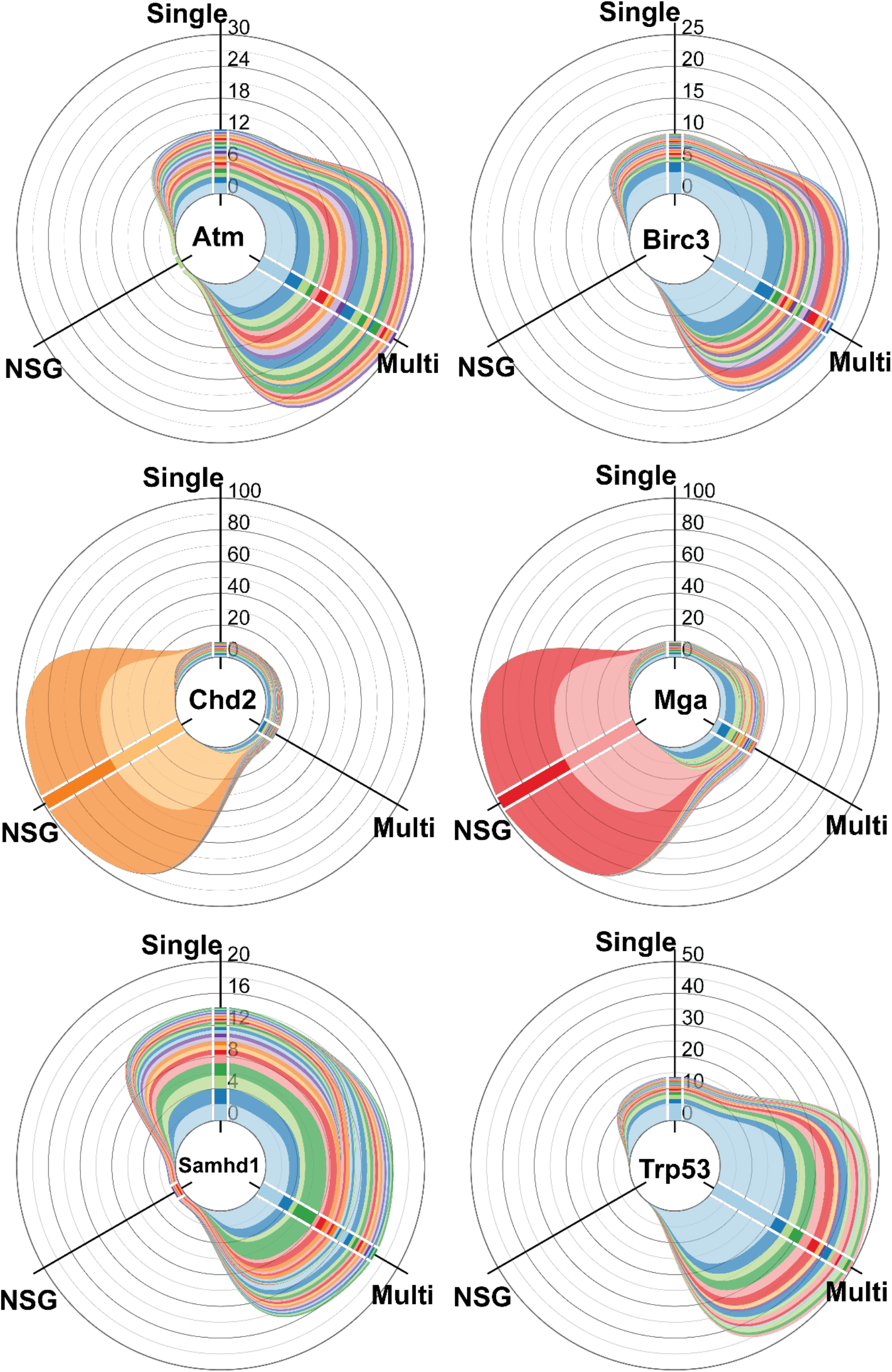
Indel profiles shared across in vitro experiments but not in vivo. Petal plots showing prevalence in percentage of cells the indels within experiment (stacked bar plots at each axis) and its prevalence in other experiment (colored bands connecting stacked bar plots). Three axes represent three experiments, single edit (Single), multi edit (Multi), and NSG. Each colored box in the stacked bar plot on the axis represents an indel sequence and its thickness corresponds to the percentage of cells the indel sequence is detected. Color matched ribbons/bands connecting across the axis are used to show that same indels found in different experiments. There are 6 plots one for each target locus.

### Challenges of translating reads count to abundance in cell population

Thus far indel profiles for each experiment provided insight into the distribution of the indels at cell population and the allelic editing difference in individual cell, the homo or hetero zygosity. These cell level details are lost in bulk analysis masking the true estimate of edited cell population as well as the zygosity of the population. It remains unclear extent of the under or over estimation of the editing in cell populations and single cell editing can be used to address this issue. Positive correlation (Pearson’s r) between the read count of indel sequences (% reads) and cell abundance (% cell with the indel) indicates a linear relation between read percentage and abundance of indel in cell population (Figure 5). However, since the correlation is not consistent across all the target loci interpretability of this relation will vary. Further, the low R2 value of the regression analysis indicates that estimation of the edited cells using read frequency is less than ideal. Data from NSG mouse was excluded due to presence of very few indel sequences and dominance of 4 indel sequences of two target gene loci. Since in >95% cells scEDIT identified heterozygous editing for Chd2 and Mga however in bulk analysis these details will be hidden. The correlation and regression analysis shows that the read count can provide a reasonable estimate of level of editing in cell population. However, read count is inadequate measure to quantitate true editing level and importantly the zygosity of the edited cell population. Single cell analysis overcomes these critical limitations of bulk cell analysis and provides insight into the indel profiles at single cell resolution. scEDIT can identify and quantify editing in massively parallel base editing at single cell level: Martin-Rufino et al. transduced human HSPCs with a pooled gRNA library, followed by electroporation with base editors (ABE or CBE). The cells were either maintained or induced to differentiate into specific blood lineages, such as erythroid cells. gRNAs with ABE were designed to introduce edits in two genes: GATA1, a master transcription factor essential for erythroid and megakaryocyte lineage differentiation, and the regulatory region of HbF, which is associated with hemoglobinopathies. In addition to single-cell RNA-seq, pooled single-cell targeted DNA sequencing was performed using the Tapestri platform.

**Figure 5.**
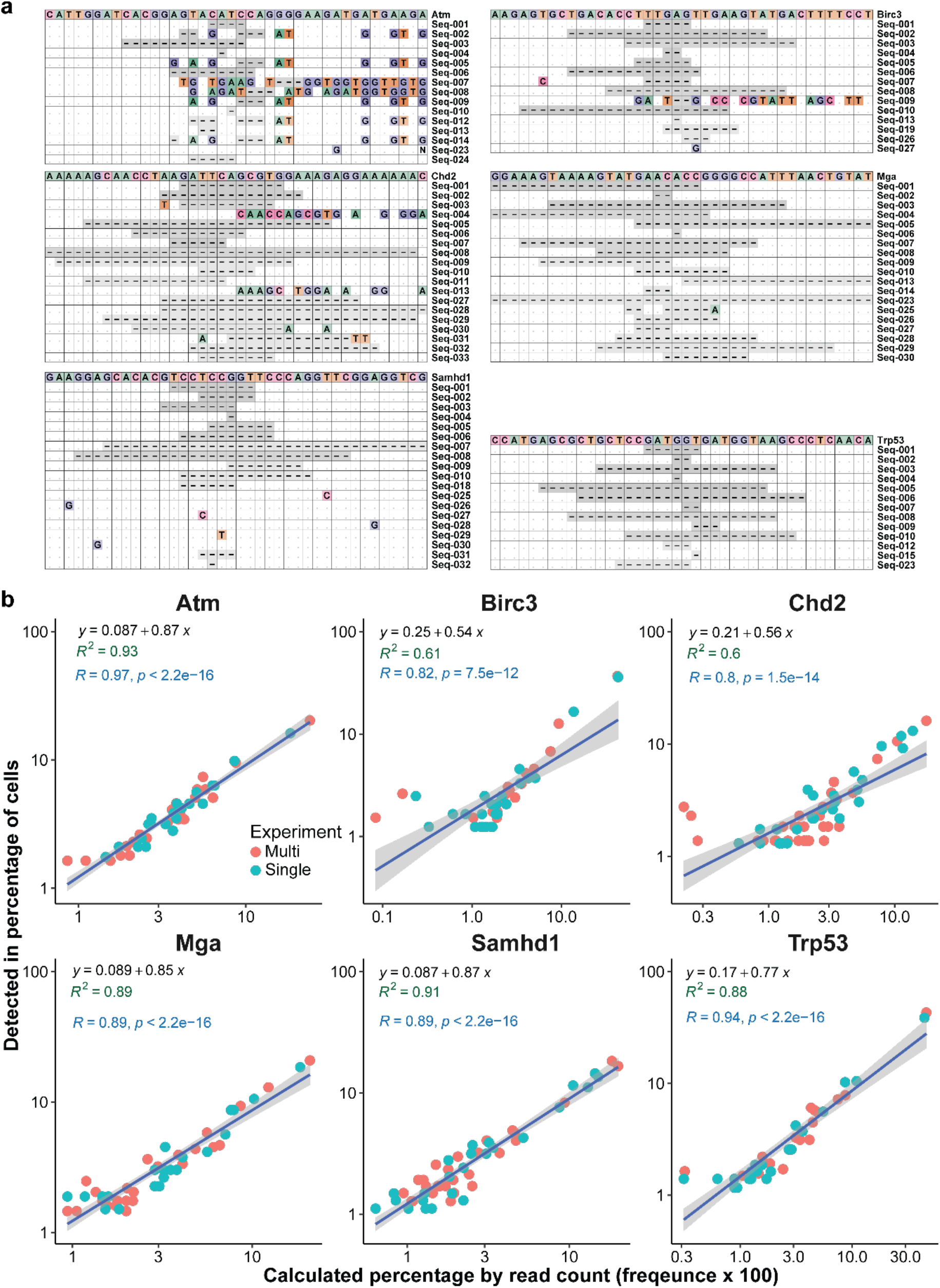
Abundance of indel sequence by read count is proportional to its prevalence in the cell population. a) Indel profiles of 6 target loci with indels ordered by their prevalence in the cell population. In indel profile plots the topmost sequence is reference sequence, mismatched bases are shown with respective nucleotides and deletion, and match are shown using ‘-’ and ‘.’, respectively. b) Scatter plot for each target locus shows the correlation between indel frequency (percentage =100 x frequency) by read count and abundance by cell count. Each colored dot represents frequency and abundance of a unique indel sequence. Regression lines are shown for two experiments with predicted equation (in black text), R-squared value (Green text), and correlation parameters Pearson’s R and associated p-value (in blue text).

To analyze this data, scEDIT is run in the BaseEdit mode to identify cell barcodes and gRNA integrated in the cells and then detects the editing outcome in all amplicon sequences. Martin-Rufino et al. performed four replicates for GATA1 and two replicates for HbF edit experiments. However, due to high data volume only partial data from each HbF replicates was used for the analysis. In this study, scEDIT efficiently processed the entire large datasets on a desktop computer.

Downstream analysis of the scEDIT-generated data determined cell identity by both cell barcode and gRNA sequence detected in the cell. Only edits within the cell-associated gRNA edit window were considered true edits for the final determination of editing status, while edits outside the edit window of the cell-specific gRNA were disregarded. Analysis revealed the distribution of gRNA and the edited cell population. In four replicates of the GATA1 experiment, the percentage of edited cells ranged from 6.6% to 26.3%, showing noticeable variability compared to that observed between two replicates of the HbF experiments (17.3% and 18.1%) (Figure 6A). Although editing patterns in most edited cells indicated homozygous editing, data revealed the presence of heterozygous editing, with two alleles having a different number of edits and/or locations within the edit window. In HbF targeting experiments, the heterozygous cell population is higher than the homozygous population, and this heterozygous editing is also higher compared to that in the GATA1 study (Figure 6A). Further analysis of gRNA at the single-cell level revealed that some gRNAs are either not present or may be below detection level. Analysis of gRNA abundance revealed a pronounced unevenness in their distribution across the cell population. A small subset of gRNAs exhibited high abundance, appearing in numerous cells, whereas the majority of gRNAs were detected in only a limited number of cells (Figure 6B). Analysis of the total number of cells in which a gRNA is detected versus the percentage of edited cells shows a positive correlation (Spearman rho = 0.54); however, with an R-squared value of 0.38 (regression analysis), it remains unclear if editing efficiency is linearly related to gRNA dominance (Supplementary Figure 3). These results demonstrate that massively parallel single-cell assays can provide a cell population-level distribution of hundreds of gRNAs, their edit zygosity, and editing efficacy. This single-cell DNA-seq assay can also provide sequence-level editing activity of these gRNAs at single-cell resolution.

**Figure 6.**
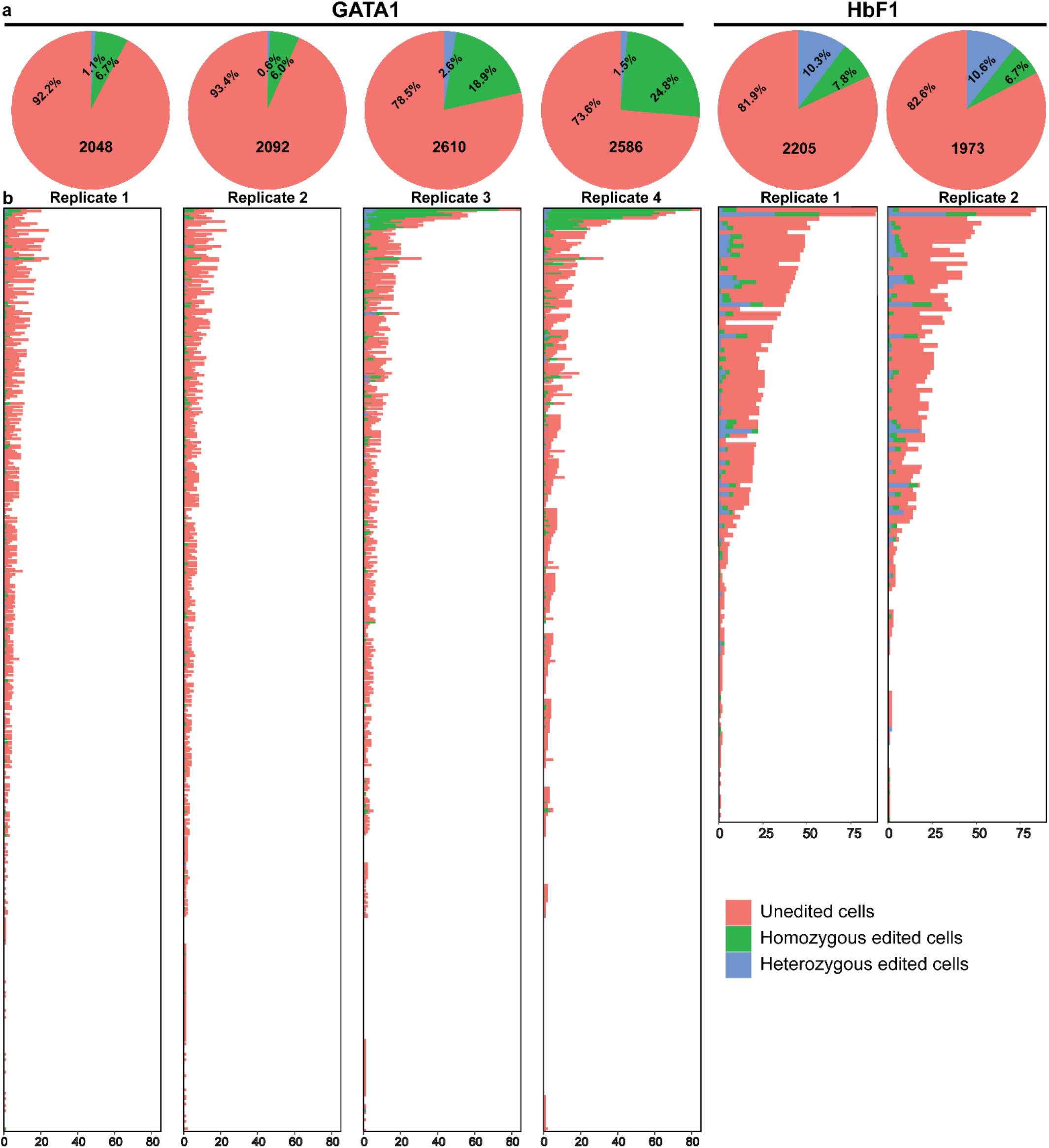
scEDIT estimate of number of cells identified for each replicate and proportion of base edited cells for gRNA detected per cell. a) Pi-charts showing the percentage of cells unedited, heterozygous, and homozygous edited cells for four GATA1 replicates and two HbF replicates. b) Barplots showing distribution of gRNAs across unedited, homozygous and homozygous edited cell populations for four GATA1 replicates and two HbF replicates.

#### Base editing biased towards start and end of gRNA edit window

The single cell data analysis of hundreds of gRNA showed skewed dispersion of gRNA across cell population. The advantage of single cell DNA-seq is the ability to explore the sequence level outcomes the gRNA and when combined with massively parallel assay it allows for assessment of hundreds of gRNA at single cell resolution. For improved accuracy the analysis editing activity of the dominant gRNA is further investigated. For GATA1 top 50 gRNAs by the cell count from 4 replicates are considered for further analysis (Figure 7A bar plot). The data of these gRNAs from both edited and unedited sequences of gRNA associates cell used to estimate editing frequency at each nucleotide within the editing windows (Figure 7A). In GATA1 experiments, top gRNAs also showed high proportion of edited cells with high single site editing frequency. For some gRNAs edits are observed different locations within the edit window. Editing of multiple nucleotides in the same cell were also observed. Majority of these edits are A>G consistent with the use of ABE used in the GATA1 and HbF experiments and T>C are the second most common edits (*11*). Substitutions and deletions are also detected, however, at much lower frequency compared to base edits.

**Figure 7.**
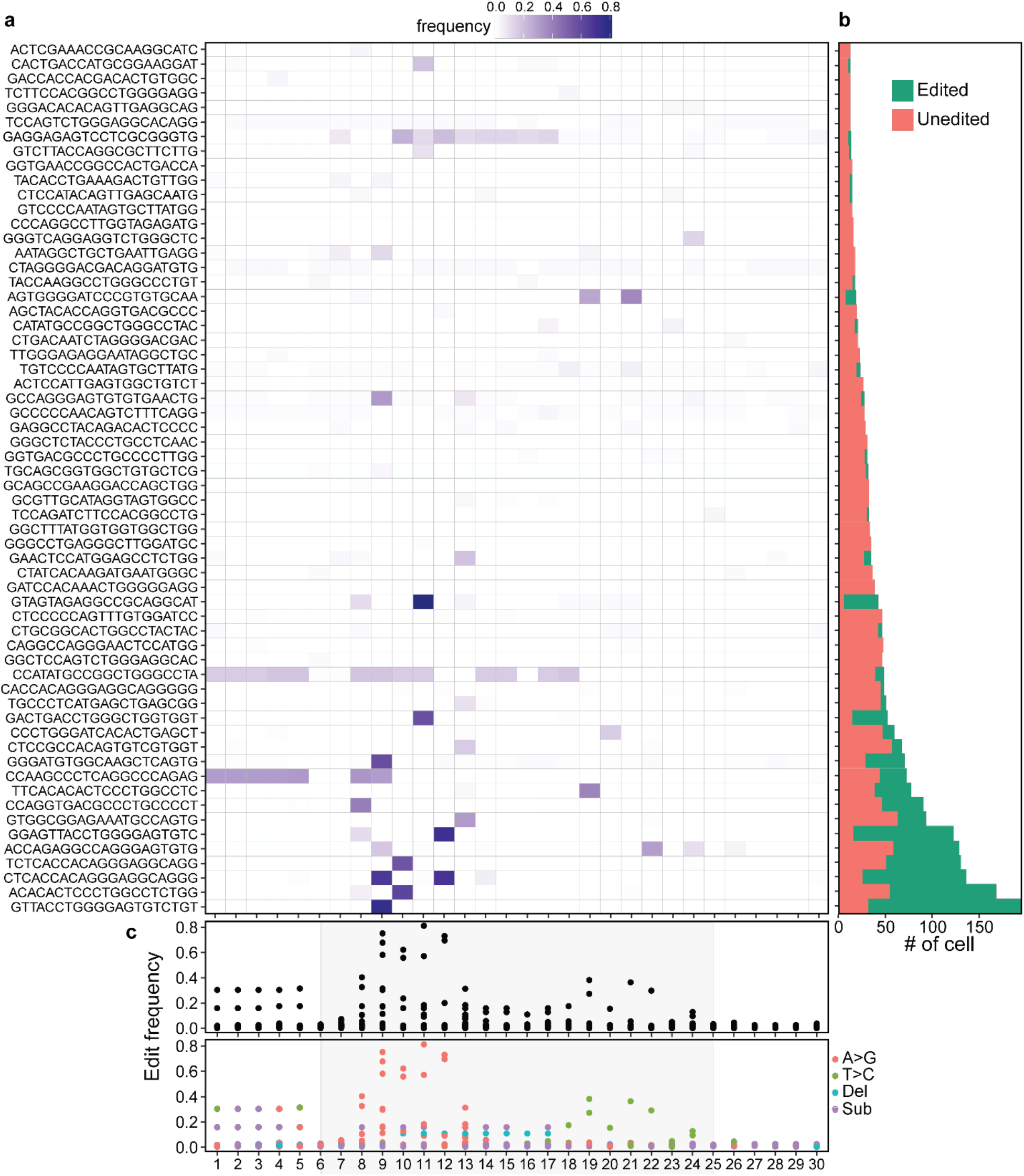
Editing patterns for GATA1 targeting gRNAs. a) Editing patterns of top 50 gRNAs by cell count from each of the four replicates. Tile plot showing editing frequence at each base in editing window (30base window: 5 bases upstream and downstream of 20 base gRNA). b) Bar plot showing total number of edit and unedited cells (summed over all replicates) associated with the top gRNAs. c) Plot showing overall and specific editing frequency at each base in the editi window (gray shaded region marks gRNA position).

For the HbF experiments, data shows similar editing patterns as observed in the GATA1 experiment (Figure 8). Key differences arise from the editing efficiency of gRNA targeting HbF seen as lower percentage of edited cells for each gRNAs (Figure 8 bar plot). Similar to GATA1 experiments, use of ABEs also led to higher A>G editing compared to T>C. Substitutions events are also observed however deletion events are more frequent than that observed in GATA1. Data from both the experiments indicated the propensity to edit sequences at the start and end of the gRNA sequence (Figure 9). A>G edits are more prevalent at the start and T>C towards ends of the gRNA an observation consistent with the other studies. Importantly, analysis revealed higher than expected rate of unintended deletion events. The single cell analysis has the advantage that greatly reduces the dilution of reads that occurs in bulk cell analysis. Such unintended deletion can be detrimental when considering ABEs for therapeutic purposes and single cell data provides a unique opportunity in detecting such rare events. Additionally, with single cell data activity of gRNA in each cell represents a unique edit event or sample point. Measuring gRNA activity in multiple cells in same experiment would result in single cell replicates and the data from multiple single events can be used for optimizations of ABEs and gRNA design.

**Figure 8.**
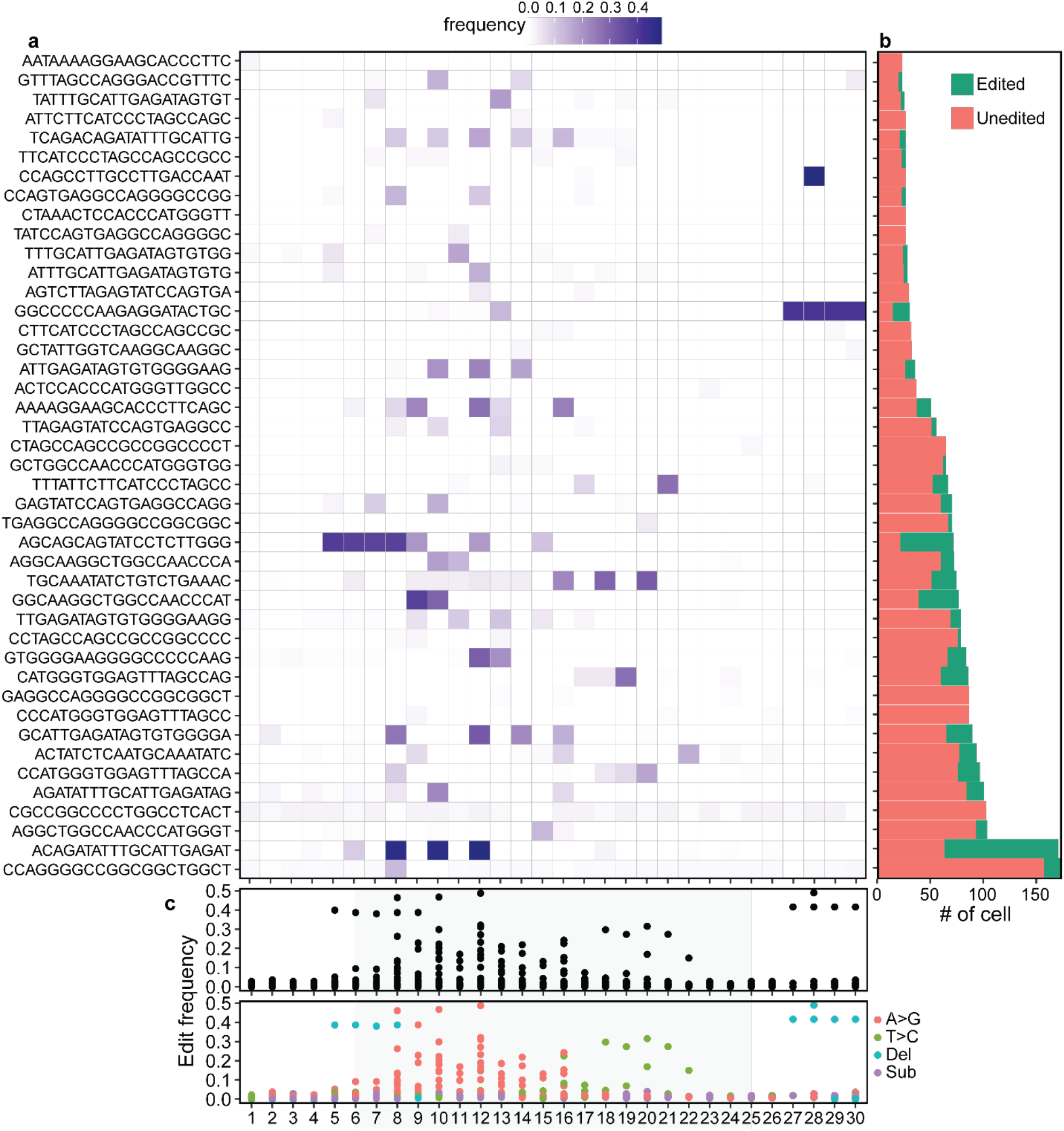
Editing patterns for HbF targeting gRNAs. a) Editing patterns of top 75 gRNAs by cell count from each of the four replicates. Tile plot showing editing frequence at each base in editing window (30base window: 5 bases upstream and downstream of 20 base gRNA). b) Barplot showing total number of edit and unedited cells (summed over all replicates) associated with the top gRNAs. c) Plot showing overall (top plot) and frequency specific edits (bottom plot) at each base in the editing window (gray shaded region marks gRNA position). Each point at a base position represents a frequency of overall or specific edit frequency for a unique top gRNA at that base position.

**Figure 9.**
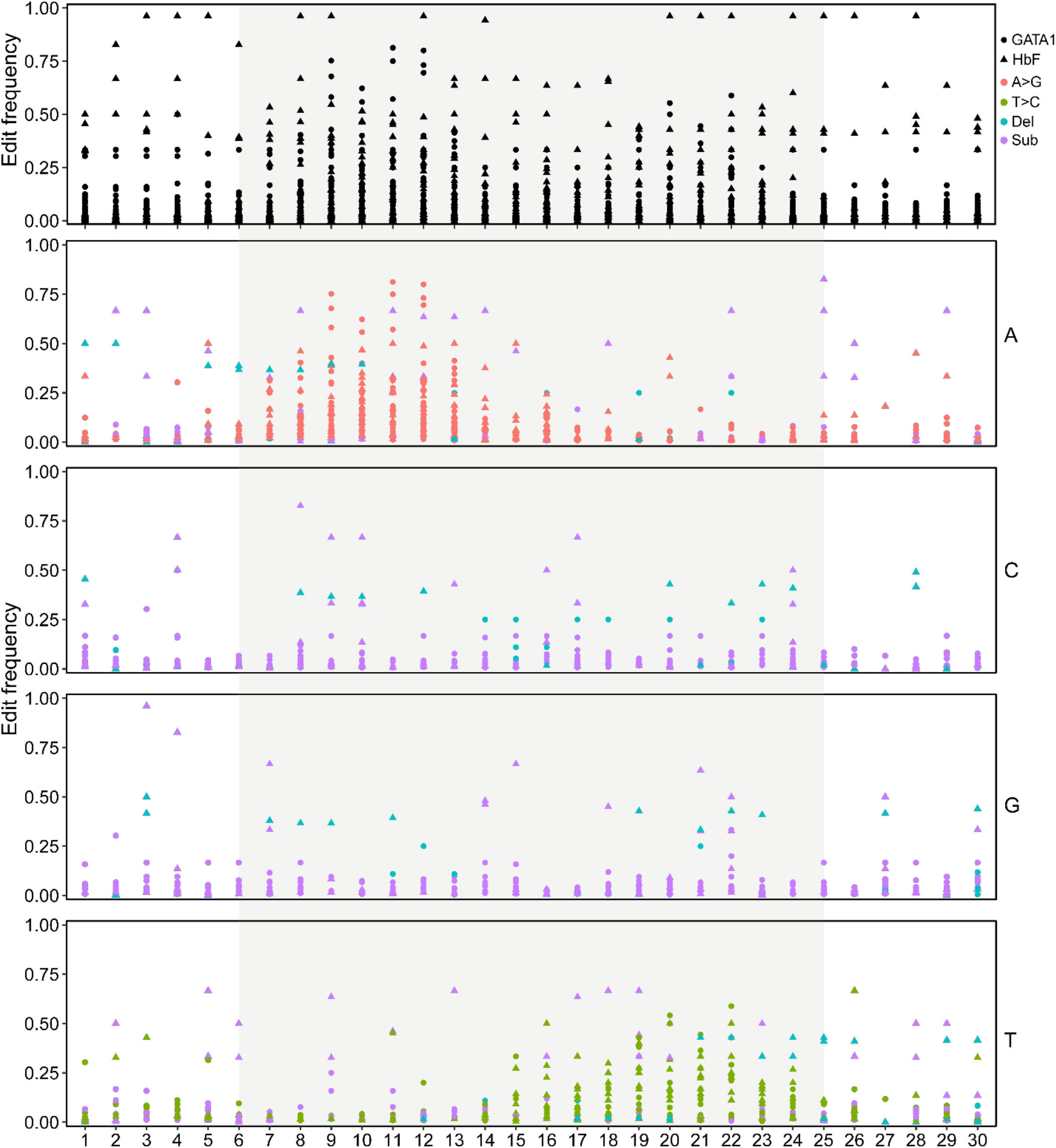
Overall editing pattern showed bias for start and end of gRNA. Black dots of the topmost plot show the overall editing frequency at each base in the editing window for all the gRNAs from GATA1 (round) and HbF (triangle) experiments. Bottom four plots show the editing frequency for a given nucleotide (A, C, G, and T shown on right side of the plot) at each location within the editing window. Different shapes of are used to represent GATA1 (round) and HbF (triangle). Colors are used to indicate type of edit outcome and gray shaded region marks gRNA position.

#### Using single cell data for machine learning to predict editing outcomes of gRNA

For GATA1 and HbF1 experiments, scEDIT provides ∼16,000 observations from 13,514 single cells (9,336 cells from 4 GATA1 replicates and 4,178 cells from 2 HbF1 replicates) of 651 gRNA sequences. Each edit event from single cell represents a unique independent experiment thus each single cell experiment provides broad range of variability of multiple gRNAs in one experiment. With bulk cell experiments of gRNA editing, outcomes are diluted by the unedited sequences from cells with gRNA failed to edit. Therefore, single cell editing studies provide more reliable data on variability of edit outcomes and reduce dilution of rare edit events. To devise a machine learning model for gRNA optimization using experiments, Doench et al one hot encoding (*12*) was used to convert 15,929 observations for 651 gRNA to numerical data. Three different classifiers support vector machine (SVM), random forest, and extreme gradient boosting were tested to model the single cell data. gRNAs were randomly split 80:20 to separate the observations into train and test data sets and all three models were trained using the training data set. Results of model testing revealed that all three models had similar accuracy of ∼0.79, however the AUC scores indicate that both random forest and extreme gradient boosting performed better than SVM. These machine learning models also provide features of importance, demonstrating the utility of scEDIT data for optimization of gRNA for improved activity. These results show that massively parallel assays in combination with single cell sequencing provide critical data for developing efficient and safe gRNAs.

## Discussion

Single cell sequencing is becoming a preferred method to study the heterogeneity of cell population. Single cell RNA-seq has become an essential tool for exploring the transcriptomic level heterogeneity and for drug discovery. The single cell DNA sequencing platform such as Tapestri can provide single cell map of mutation landscape as well as a method to assess efficacy of gene modification introduced by vector integration or CRISPR-Cas activity. As the single cell methods are becoming affordable, reliable with high throughput capability they are likely to replace traditional bulk cell analysis. However, computational analysis of single cell data is costly in part due to the high costs of computational infrastructure required to process such large data sets. scEDIT provides a lightweight, portable, cost and time efficient for analysis of single cell DNA-seq data from CRISPR edited cells. scEDIT can efficiently run on a low-cost desktop computer and generates output that is essential for critical analysis of CRISPR edited cell population.

scEDIT demonstrated fast and efficient processing of the single cell CRISPR raw sequence data and identified valid single cells using per cell read count. Within the identified cells scEDIT can designate edited cells by detecting edits in target sequence and identifying indel profiles in each edited cell. Results from analysis differ slightly from Hacken et al study likely due to differences barcode identification and cut-offs used such as total per cells read count for cell calling, per cell amplicon read count and number of reads edited to define true editing. Since the proprietary Tapestri bioinformatics pipeline is not open-source, underlying details of algorithms used for cell calling and edit determination are only partly disclosed. Due to these restrictions a direct performance comparison of scEDIT against Tapestri bioinformatics pipeline was not possible. However, single cell analysis of CRISPR editing using scEDIT revealed indel patterns at single cell level for each of the target loci. The indel profiles are shared between the cells and the high frequency indel sequences were found in more cells and had heterozygosity with other indel sequences. This quantitative analysis of the data revealed distribution of indels in cell population. Furthermore, similar indel profile and indel distribution were observed in in vitro studies indicate presence of selection bias for certain indels over others. Interestingly, analysis of in vivo studies revealed selection bias for co-editing in both alleles of Chd2 and Mga with over 90% edited cells heterozygous editing at the two loci. Compared to common transient Cas-9 gRNA activity obtained by transfection of cells, Hacken et al used lenti-viral vector-based approach providing a sustained Cas9-gRNA activity. Lenti-viral mediated Cas-9 gRNA expression led to repeated editing activity at the loci creating to large indel(*13*). Results show that scEDIT can capture both the small and large indels introduced by the sustained Cas9 activity.

BaseEdit mode further demonstrated utility of scEDIT workflow for analysis of complex multi gRNA experiments. In the original study authors only processed partial data due to data volume limit of the Tapestri workflow. Notably, scEDIT could process analyze the full data sets on small desktop indicating its efficiency and cost effectiveness. The base editing data reveals several aspects of the Cas9-ABE activity such as editing site bias, single and multi-base editing, and rare yet critical deletions events. Additionally, data showed that integrated copy of gRNA does not result in editing activity. Causes for ineffective or lack of editing activity remain unclear however one possible explanation can be the limited bioavailability of the Cas9-ABE complex or expression level of gRNA. With ∼600 gRNA sequences single experiment was able to deliver ∼16,000 data points to initiate development of a predictive model. Larger data sets with more test gRNAs are needed to establish more accurate machine learning models with improved predictive power. scEDIT provides processed data in two formats: (1) in table format amplicon read counts for each barcode and (2) standard fastq files of filtered read sequences with each sequence name containing cell barcode and amplicon identification. Counts table format is useful for downstream analysis such cell calling, identification of edited cells and their zygosity. The fastq sequence data with cell barcode and amplicon name enables further separation of sequence by cell barcode or amplicon name. The versatile functionality of the scEDIT makes it an invaluable tool for analysis of CRISPR editing in single cell DNA sequencing data generated using Tapestri platform. Although a desktop computer is sufficient to run scEDIT, utilizing more CPU cores can significantly improve the performance thus running it on a HPC will furtherboost data processing speed. The scEDIT algorithm and script of cell barcode and amplicon detection can be utilized to process other single cell DNA-seq data set from Tapestri. Thus, scEDIT can be used as a general-purpose tool for preprocessing and as a raw count generation tool for various assays run on Tapestri platform.

## Methods

### Raw sequence processing

Raw sequence data files from Hacken et al study was downloaded from GEO () as SRA files and were converted to fastq files using SRATools. Trimmomatic (Trimmomatic-39) was used for removing Illumina adaptors and quality trimming of raw sequences. Quality trimmed sequences were then used for scEDIT input sequences. scEDIT used direct match and Smith-Waterman algorithm to locate two constant sequences present in the Tapestri forward primers. The two constant sequences are used to accurately locate and extract two 9 base pair (bp) cell barcodes separated by the constant sequences and 0-3 bp spacer. Both reads are removed if the forward read sequence is short (<25bp) or missing constant sequences. The extracted cell barcodes are then compared against whitelist of known 1536 barcodes used in combinations (1536 x 1536). scEDIT allows for maximum of 1-bp mismatch in each 9-bp barcode (total of 2-bp mismatch for total 18-bp cell barcode). Amplicons reverse and forward primers were identified by setting identity match to 90% (can be user specified and different for individual amplicon primer). Sequences missing either or both forward or reverse primers were filtered out. After identifying the amplicon primers, the sequence read was further checked for edits in the edit window, 20bp up and down of the PAM site (total 40bp sequence). First the algorithm checks for <5 base indel using SW algorithm, if no edits are detected the read is considered as unedited. In case indel short (<5 base) or single indels scEDIT check for quality score (>30) as the edit location to determine true edit from sequencing errors. Edit bases that pass the quality test are considered as true CRISPR edits. For indels longer than 5 base wavefront alignment algorithm(*14*) is used to align and identify edits in the sequences reads. scEDIT assigns each read to a cell using the barcode in the read and keeps count both edited and unedited reads for each amplicon. As a final produce scEDIT write read count data in a table format and filtered reads in fastq format.

## Post processing

scEDIT ranks cell barcodes are using the counts of associated total number of reads. Polynomial is fit to the data to determine the steep drop in total read number; this sharp bend point can be used as a cutoff point for cell calling. However, based on the experimental setup and nature of the data cutoff point can be customized for experiment specific requirements.

InDel mode: In this study, due to the nature of data the automated cell calling algorithm is unable to provide cell count. A custom cell cutoff of minimum 750 total reads/cell is used to select valid cells. Each experiment yielded in the range of 3000-3500 valid cells for the downstream analysis. BaseEdit mode:

Cell calling was done is similar fashion as for InDel mode. Base edits were called if the edit occurred inside edit-window (within 20 base gRNA target ±5 bases). scEDIT provides barcode associated read count for each amplicon including amplicons designed to capture gRNA in the integrated vector. Additionally, the tabulated data also includes information on edit status for all the gRNAs.

Valid call barcodes are used to further evaluate the level editing, heterozygosity, and to determine indel profiles. A custom R script was used for analysis of editing and zygosity in cells. A cell containing one wildtype and one indel sequence or more than 1 indel sequence indicates that the two sequences originated from two differentially edited allele. In a cell if the contribution from each allele is >20% then editing is considered as heterozygous else homozygous. In addition, for the purpose of visualizing indel profiles, a custom C++ script is used to substitute deletions with ‘-’ symbol in the original reads.

## scEDIT usage

scEDIT is available as github repository (https://github.com/GSbioinfo/scEDIT) and can be run on ubuntu and other linux distributions. In addition, a docker file to create a docker image and a sample nextflow script can be used to test and run entire scEDIT workflow. Other relevant scripts used for fastq file processing, data analysis, and indel visualization are also provided.

## Conflict of interest

The author was a visiting associate researcher at University of California, Los Angeles (UCLA). No computational resources of UCLA were used for this work. The author has no conflict of interest to declare.

## Supplemental figures

SF1: Accuracy analysis of the scEDIT. Bar plots showing percentage of read recovered by scEDIT for different target loci and reference genes.

SF2:

Indel profiles in individual cells as detected by scEDIT analysis. Figure is showing indel profiles from randomly selected few cells from two experiments. In indel profile plots the topmost sequence is reference sequence, mismatched bases are shown with respective nucleotides and deletion, and match are shown using ‘-’ and ‘.’, respectively. Indel plot titles provide in information about cell barcode (CB), experiment, read direction (R1-forward and R2-reverse), Target locus (gene) name as follows CB_EXPERIMENTRX_Target. Sequence labels show sequence #, read count, and percentage of reads with the target specific reads associated with the cell barcode. Indel sequences are ordered by percentage high to top and low to the bottom. Indel sequence with <1% reads were filtered out for visualization purposes.

SF3:

Correlation between gRNA abundance and editing outcome. Each point is unique gRNA with x-axis representing number of cell the gRNA was detected, and y-axis represents percentage of the cells where editing activity was detected within the editing window.

## Notes

### Competing Interest Statement

The authors have declared no competing interest.

### Summary of Updates

New manuscript include new feature of base edit analysis to the scEDIT software tool. Manuscript includes analysis of single cell base editing data.

