## Supplementary figures and images for "Single cell Edit Detection and Identification Tool (scEDIT): computational workflow for efficient and economical single cell analysis of CRISPR edited cells"

### Supplemental Figure 3

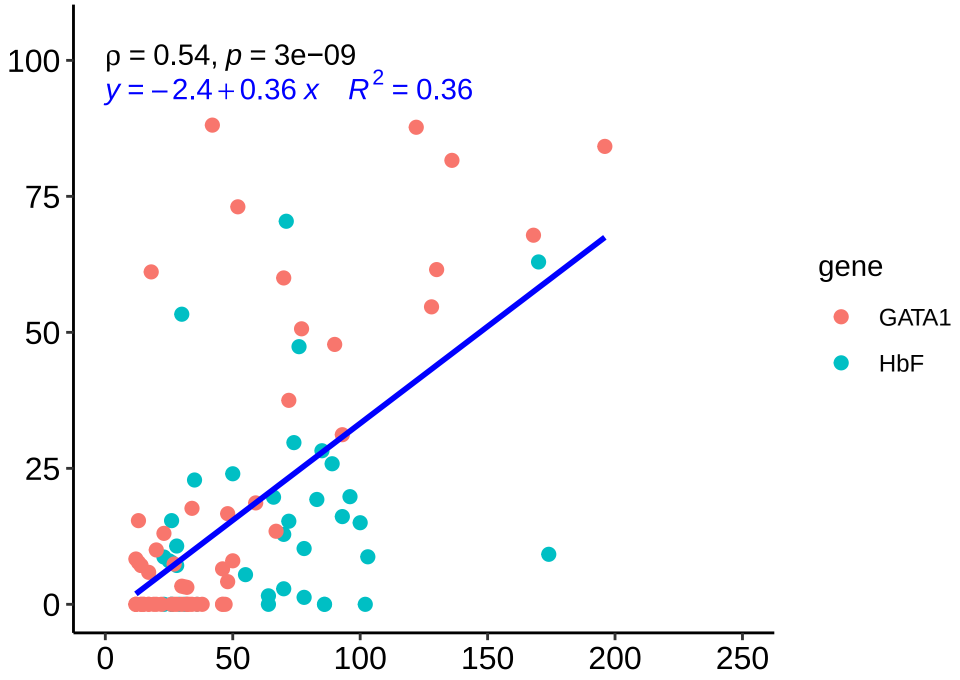
